# Geosmin attracts *Aedes aegypti* mosquitoes to oviposition sites

**DOI:** 10.1101/598698

**Authors:** Nadia Melo, Gabriella H. Wolff, Andre Luis Costa-da-Silva, Robert Arribas, Merybeth Fernandez Triana, Muriel Gugger, Jeffrey A. Riffell, Matthew DeGennaro, Marcus C. Stensmyr

## Abstract

Geosmin is one of the most recognizable and common microbial smells on the planet. Some insects, like mosquitoes, require microbial-rich environments for their progeny, whereas for other insects such microbes may prove dangerous. In the vinegar fly *Drosophila melanogaster*, geosmin is decoded in a remarkably precise fashion and induces aversion, presumably signaling the presence of harmful microbes. We have here investigated the effect of geosmin on the behavior of the yellow fever mosquito *Aedes aegypti*. In contrast to flies, geosmin is not aversive in mosquitoes but stimulates egg-laying site selection. Female mosquitoes could associate geosmin with microbes, including cyanobacteria consumed by larvae, who also find geosmin – as well as geosmin producing cyanobacteria – attractive. Using *in vivo* multiphoton imaging from mosquitoes with pan-neural expression of the calcium reporter GCaMP6s, we show that *Ae. aegypti* code geosmin in a similar fashion to flies, i.e. with extreme sensitivity and with a high degree of selectivity. We further demonstrate that geosmin can be used as bait under field conditions, and finally we show that geosmin, which is both expensive and difficult to obtain, can be substituted by beetroot peel extract, providing a cheap and viable mean of mosquito control and surveillance in developing countries.

## INTRODUCTION

The yellow fever mosquito *Aedes aegypti* is one of the world’s deadliest disease vectors, and a global health threat through the transmission of – apart from the eponymous disease – viruses responsible for dengue fever, chikungunya, and Zika. Much effort has accordingly been spent on deciphering how *Ae. aegypti* (and other mosquitoes) locates human hosts [reviewed in Montell and Zweibel, 2016], with the aim of devising means to prevent this from happening [E.g. USDA, 1947]. Another approach to prevent pathogen transmission is to disrupt the reproductive cycle of the mosquitoes. For example, targeting the oviposition stage of the female mosquito would reduce vector density and hence decrease the chances of future epidemics [Barrera *et al*., 2014; 2017].

Following a successful blood meal, the female mosquito sets off in search of a suitable egg-laying site, such as stagnant water [Bentley and Day, 1989]. To find oviposition sites, mosquitoes rely on a combination of hygrosensation and olfaction, with the latter used to sense volatiles produced by aquatic microbes, which together with plant detritus serves as food for the larvae [Bentley and Day, 1989]. Microbes generate a plethora of volatile chemicals, of which several has been shown to confer oviposition site selection in mosquitoes [e.g. Ponnusamy *et al*., 2008; 2011], whereas others induce avoidance [Huang *et al*., 2006]. Microbial volatiles can accordingly be used to manipulate oviposition behavior in mosquitoes.

Geosmin is a volatile compound produced by a wide range of microorganisms, including taxa that inhabit typical mosquito breeding sites [Wang *et al*., 2011; Vasquez-Martinez *et al*., 2002]. To the human nose, this chemical has a rather pleasant and an immediately recognizable smell of wet soil (**Figure 1A**)(or beetroot, pending on your cultural background). To the vinegar fly *Drosophila melanogaster*, however, geosmin signals the presence of harmful microbes and is innately aversive [Stensmyr *et al*., 2012]. Interestingly, the olfactory system of both humans and flies are extremely sensitive to geosmin [Polak and Provasi, 1992; Stensmyr *et al*., 2012], with flies even equipped with a functionally segregated olfactory channel exclusively mediating information regarding this chemical [Stensmyr *et al*., 2012]. How mosquitoes perceive this important and characteristic microbial smell remains, however, unknown. We have here investigated the effect of geosmin on *Ae. aegypti*. In stark contrast to flies, geosmin is not aversive to *Ae. aegypti*, instead it stimulates egg-laying site selection. Gravid female mosquitoes likely associate geosmin with aquatic microbes, including cyanobacteria, which are consumed by larvae, who also find geosmin – as well as geosmin producing cyanobacteria – attractive. Using *in vivo* two-photon imaging from mosquitoes with pan-neural expression of the calcium reporter GCaMP6s, we show that *Ae. aegypti* code geosmin in a similar fashion to flies, i.e. with extreme sensitivity and with a high degree of selectivity. We further demonstrate that geosmin can be used in the field as bait to lure female mosquitoes. Finally we demonstrate that geosmin, which is both expensive and difficult to obtain, can be substituted by beetroot peel extract.

**Figure 1.**
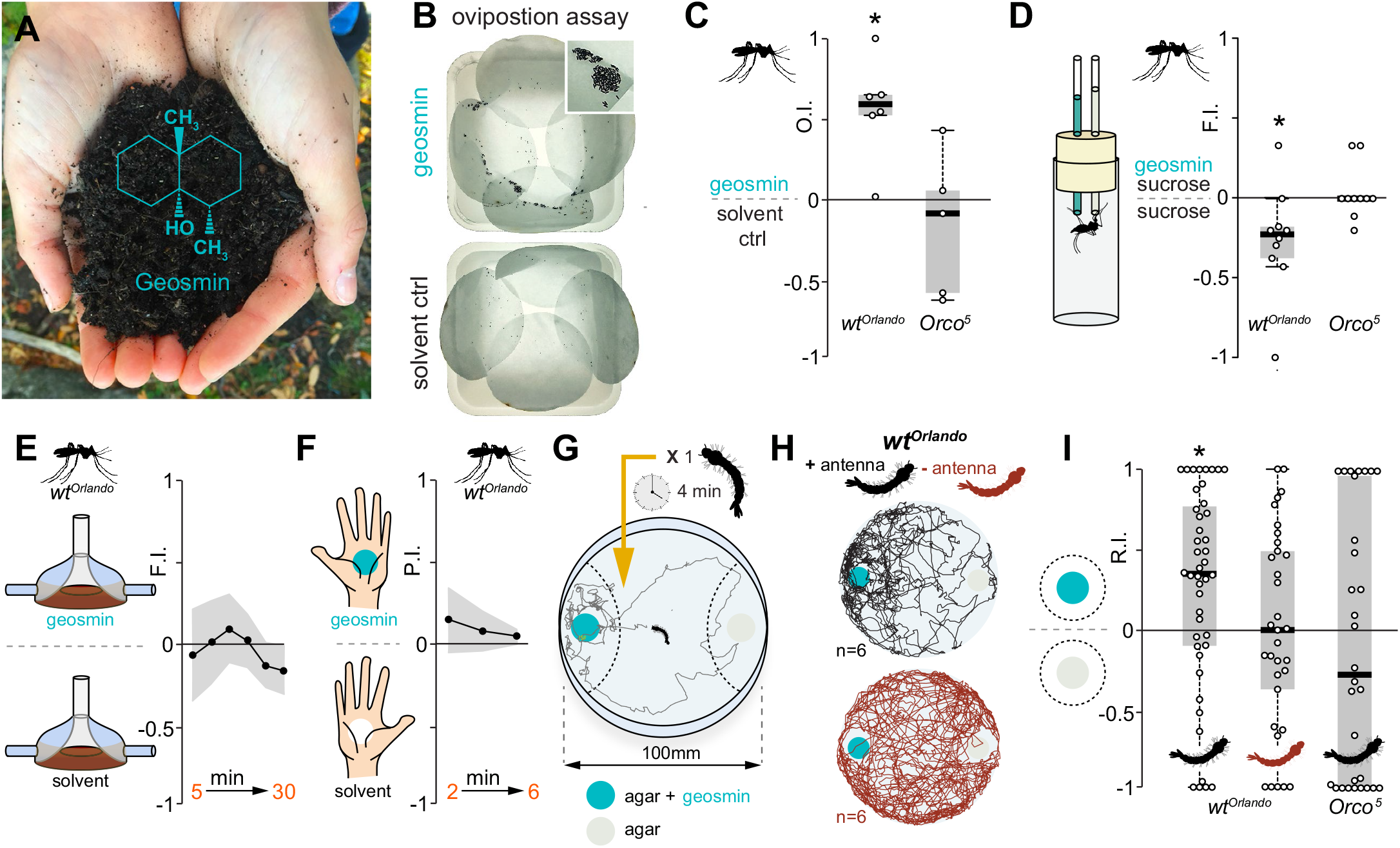
Geosmin confers egg-laying selection in *Aedes aegypti*. (**A**) Geosmin has to the human nose the distinct smell of wet soil, and is produced by a wide range of microorganisms, both terrestrial and aquatic. Photo: M. Stensmyr (**B**) Plastic trays lined with filter paper used in oviposition experiments. Top, water with geosmin added, below, water only control. Insert: close-up of a cluster of *Aedes* eggs in the geosmin containing tray. (**C**) Oviposition indices (OI) of WT (20 mosquitoes/trial, n=6) and *Orco^5^* mosquitoes (20 mosquitoes/trial, n=6 trials) from a binary-choice test between water and water spiked with geosmin (10^−5^). The edges of the boxes are the first and third quartiles, thick lines mark the medians, and whiskers represent data range. Preference was tested with one-sample Wilcoxon test, theoretical mean 0. Star denotes significantly different from 0; p < 0.05. (**D**) Feeding indices (FI) from a CAFE assay of WT (n=10) and *Orco^5^* mosquitoes (n=10) given a choice to feed from two capillaries with succrose water (10%), once of which in addition containing geosmin (10^−3^). Preference was tested with one-sample Wilcoxon test, theoretical mean 0. Star denotes significantly different from 0; p < 0.05. (**E**) FI over 30 min from WT mosquitoes (20 mosquitoes/trial, n=5) given a choice to feed from two membrane blood feeders, one of which scented with geosmin (10^−5^). Shaded line indicates SEM. Preference was tested with one-sample Wilcoxon test, theoretical mean 0, for each time point. (**F**) Probing index (PI) from WT mosquitoes (20 mosquitoes/trial, n=5) in a constrained contact assay over 6 min, provided with a choice to approach and probe two hands (from the same individual), one of which scented with geosmin (10^−3^). Shaded line indicates SEM. Preference was tested with one-sample Wilcoxon test, theoretical mean 0, for each time point. (**G**) Schematic of the larval behavioral assay. Dashed lines denote the two zones in which time spent was measured. (**H**) Sample tracks of WT larvae with antennae (above) and with antennae removed (below). (**I**) Response indices of WT larvae with antennae (n=44), without antennae (n=33), and *Orco^5^* mutants (n=30). Box plots as per (**C**). Preference was tested with one-sample Wilcoxon test, theoretical mean 0. Star denotes significantly different from 0; p < 0.05.

## RESULTS AND DISCUSSION

### Geosmin confers oviposition site selection and larval attraction

In flies, geosmin signals the presence of harmful microbial contaminants and negatively affects egg-laying preference [Stensmyr *et al*., 2012]. We similarly examined if geosmin also affects egg-laying preference in the mosquito. Female mosquitoes provided with a choice to oviposit in containers with water, or water spiked with geosmin (10^−5^ dilution), strongly preferred to lay eggs in the latter (**Figure 1B**). Thus, in contrast to *D. melanogaster, Ae. aegypti* evidently perceives geosmin as attractive. *Ae. aegypti*, like other insects, detects odors via members of two large gene families; odorant receptors (ORs)[Vosshall *et al*., 1999; Clyne *et al*., 1999] and ionotropic receptors (IRs)[Benton *et al*., 2009]. The egg-laying preference towards geosmin is mediated by the olfactory system, since assays with *Orco^5^* mutants [DeGenaro *et al*., 2013] – a co-receptor needed for proper OR function [Larsson *et al*., 2004] – revealed no difference in egg numbers between water and water treated with geosmin (**Figure 1C**).

Other behaviors, however, were barely or only moderately affected by the presence of geosmin. Mosquitoes presented with a choice of sucrose water (10%) versus sucrose water mixed with geosmin (10^−3^) in a CAFE assay [Ja *et al*., 2007] (**Figure 1D**) – an assay that here would largely mimic nectar feeding – showed a slight aversion to feeding from the geosmin scented capillaries (**Figure 1D**), which was not observed in *Orco^5^* mosquitoes (**Figure 1D**). Addition of geosmin (10^−5^) to a membrane blood feeder revealed no adverse effect on blood feeding compared to unscented feeders (**Figure 1E**), and similarly, addition of geosmin (10^−3^) in a constrained contact assay showed also no negative effects on host attraction (**Figure 1F**)(See Experimental procedures).

We next asked how *Aedes* larvae react to the presence of geosmin in their aquatic habitat. To address this issue, we devised a larval two-choice assay, which allowed us to monitor the position of single larvae over time (**Figure 1G**). 3^rd^ and 4^th^ instar *Ae. aegypti* larvae showed positive chemotaxis towards geosmin, although with considerable individual variation (**Figure 1H, I**). As with the adults, this behavior was dependent upon olfaction, since larvae with ablated antenna showed no preference (**Figure 1H, I**), and moreover, dependent upon the activation of *Orco* positive neurons (**Figure 1I**). In summary, geosmin specifically mediates oviposition site selection in *Ae. aegypti* and olfactory guided positive chemotaxis in larvae.

### Geosmin producing cyanobacteria confer oviposition and larval attraction

A plausible assumption would be that geosmin signals the presence of microbes to *Ae. aegypti*, akin to its function in flies [Stensmyr *et al*., 2012], albeit with opposite valence. In the habitats of the aquatic larvae, cyanobacteria are a common source of geosmin, and have also been isolated from the gut of wild mosquitoes [Thiery *et al*., 1999; Vazquez-Martinez *et al*., 2002; Wang *et al*., 2011]. We first examined how adult *Ae. aegypti* reacts to cyanobacteria. We selected a potentially geosmin-producing strain, *Kamptonema* sp. PCC 6506 [Calteau *et al*., 2014], verified geosmin production via solid phase micro-extraction (SPME) and gas-chromatography mass spectroscopy (GC-MS) (**Figure 2A**), and then performed oviposition choice experiments with wildtype *Ae. aegypti* females. Water (60 mL) inoculated with cyanobacteria (250 μL of the bacterial suspension) was clearly preferred over water with only growth medium added (**Figure 2B**). This preference was dependent upon activation of *Orco* expressing neurons (**Figure 2B**).

**Figure 2.**
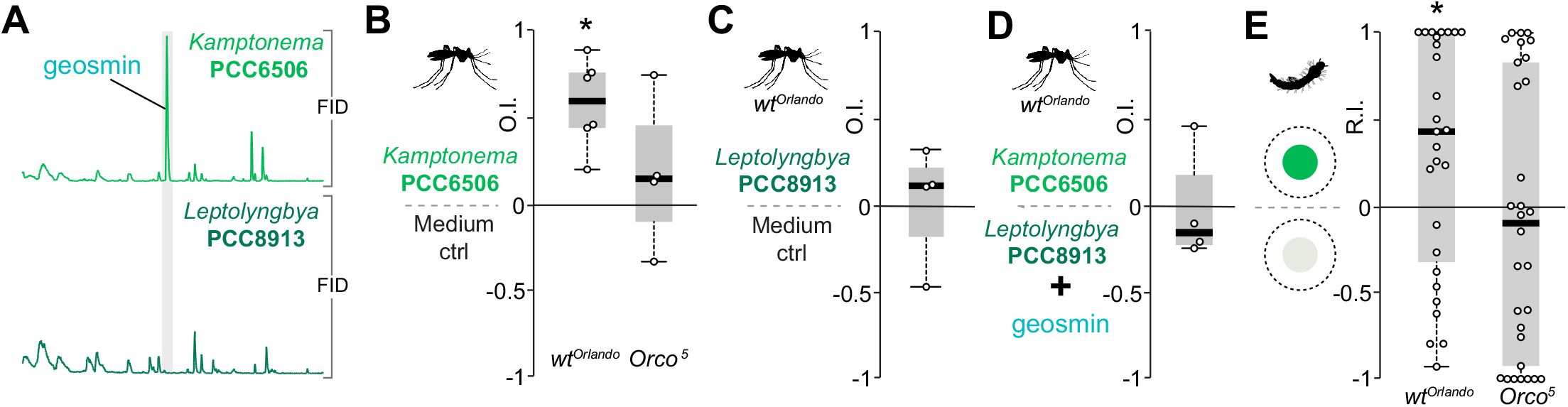
Geosmin producing cyanobacteria confer egg-laying and larval attraction. (**A**) Flame ionization detection (FID) traces from a gas chromatography-mass spectrometry analysis of head space volatiles emitted by two strains of cyanobacteria. (**B**) Oviposition indices (OI) of WT (20 mosquitoes/trial, n=6) and *Orco^5^* mosquitoes (20 mosquitoes/trial, n=4) from a binary-choice test between growth medium and growth medium with a cyanobacteria strain (PCC6506) producing geosmin. Box plots as per Figure 1C. Preference was tested with one-sample Wilcoxon test, theoretical mean 0. Star denotes significantly different from 0; p < 0.05. (**C**) OI of WT mosquitoes (20 mosquitoes/trial, n=4) from a binary-choice test between growth medium and growth medium with a cyanobacteria strain (PCC8913) not producing geosmin. Box plots as per Figure 1C. Preference was tested with one-sample Wilcoxon test, theoretical mean 0. (**D**) OI of WT mosquitoes (20 mosquitoes/trial, n=4) from a binary-choice test between PCC 6506 and PCC 8913, the latter with geosmin added. Box plots as per Figure 1C. Preference was tested with one-sample Wilcoxon test, theoretical mean 0. (**E**) Response indices from larvae (WT, n=27; *Orco^5^*, n=32) given a choice between agar mixed with growth medium and agar with PCC 6506. Box plots as per Figure 1C. Preference was tested with one-sample Wilcoxon test, theoretical mean 0. Star denotes significantly different from 0; p < 0.05.

As evident from the GC-MS profile (**Figure 2A**), *Kamptonema* sp. PCC 6506 produces in addition to geosmin, a range of other volatile chemicals, which begs the question whether or not geosmin alone mediates the preference. To address this issue, we selected another cyanobacterial strain isolated from a mosquito-breeding site (*Leptolyngbya* sp. PCC 8913)[Thiery *et al*., 1999] not producing geosmin, as verified via SPME and GC-MS (**Figure 2A**). We then performed the same oviposition choice experiments as with *Kamptonema* PCC6506. The female mosquitoes now displayed no preference for the cyanobacteria containing vessels (**Figure 2C**). We then provided the mosquitoes with a choice of *Kamptonema* PCC 6506 against *Leptolyngbya* sp. PCC 8913, with the latter spiked with geosmin (5 ng pure substance in 60 mL water), an amount roughly equivalent to the release of geosmin from PCC 6506, as determined by SPME/GC-MS (data not shown). Mosquitoes confronted with this choice, showed no preference either way (**Figure 2D**).

We next examined how larvae react to the presence of cyanobacteria. Larvae screened in the same two-choice assay as before, showed an overall preference to the side baited with *Kamptonema* sp. PCC 6506, again with quite some individual variation (**Figure 2E**). Similar to the egg-laying behavior of the adults, the larval positional preference was also dependent upon *Orco* neurons (**Figure 2E**). We conclude that geosmin is key to the preference of egg-laying in water containing cyanobacteria. This preference is also observed in larvae, which presumably associate geosmin with the presence of food.

### Two-photon imaging reveals sensitive and selective neural coding of geosmin

To examine how *Ae. aegypti* smells geosmin, we next performed electroantennography (EAG) from wild type *Ae. aegypti* (Orlando). EAGs revealed distinct base line deflections in response to stimulation with geosmin (**Figure 3A**), suggesting that the antennae house olfactory sensory neurons (OSN) tuned to this microbial volatile. In line with the oviposition experiments, EAGs from *Orco^5^* mutants [DeGennaro *et al*., 2013] showed no geosmin (or 1-octen-3-ol) induced antennal responses, whereas octanoic acid, a compound detected by the IR pathway [Prieto-Godino *et al*., 2017], induced responses no different from those obtained with the *Orlando* wildtype control.

**Figure 3.**
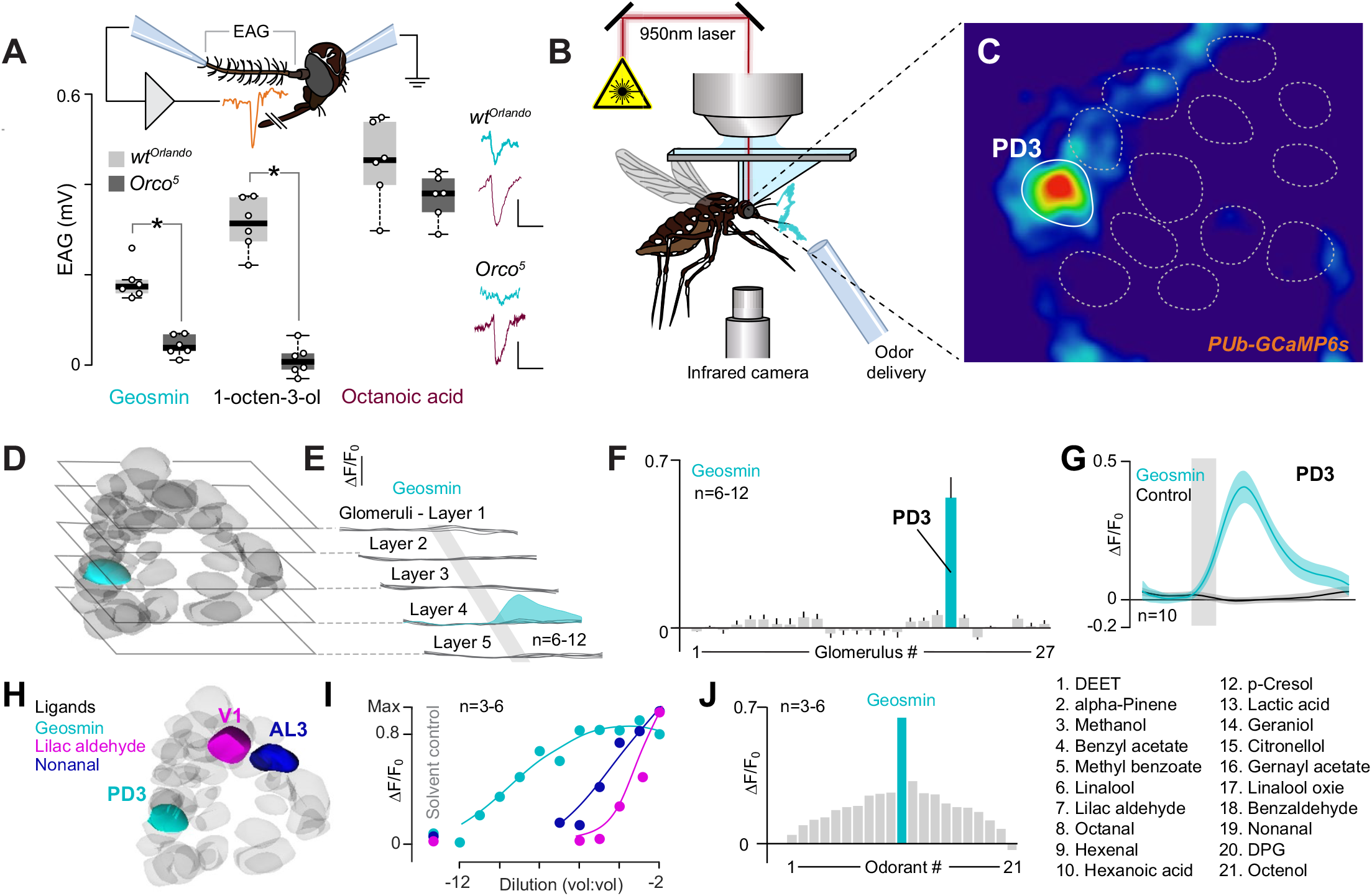
*Aedes aegypti* detects geosmin with extreme sensitivity and seletivity. (**A**) Schematic of the electroantennogram (EAG) preparation (top). EAG responses from WT and *Orco* mutants to stimulation with 10^−3^ dilutions of geosmin, 1-octen-3-ol, and octanoic acid. Right, representative recordings. Vertical scale bar 0.25 mV, horizontal bar 5 s. Statistical difference was measured via a Student’s t-test. Star denotes significant difference (p < 0.05). (**B**) Schematic of the two-photon setup used to record calcium dynamics in the mosquito antennal lobe (AL). (**C**) Pseudocolor plot of ΔF/F_0_ calcium responses (0-1 scale) to geosmin (10^−3^ dilution), at a depth of 75 μm from the surface of the AL. Geosmin evoked a strong response in one glomerular region-of-interest (highlighted in white). (**D**) Non-responsive AL glomeruli (grey) and the geosmin-responsive glomerulus (green; the third posterodorsal glomerulus [PD3]) registered and mapped to an AL atlas and cross-referenced to a previously published atlas [Ignell et al., 2005]. (**E**) Glomerular responses (ΔF/F_0_) to geosmin characterized at five depths (15, 30, 50, 75, and 90 μm) from the ventral surface of the AL. Each trace is the mean of one glomerulus; the PD3 response is shown in green. Vertical scale bar: 0.4%. Grey bar denotes stimulus duration (2 s). (**F**) Responses to geosmin across all sampled glomeruli; only the PD3 glomerulus (bar in green) showed significant calcium dynamics to geosmin compared to the solvent control (Kruskal-Wallis test: p < 0.05). Bars represent the mean ± SEM. (**G**) Dynamics of the calcium response to geosmin (green trace) and the solvent control (DPG, black trace) for the PD3 glomerulus. Lines are the mean; shaded areas are the SEM. Grey bar denotes stimulus duration (2 s). (**H**) AL atlas showing the PD3 glomerulus (green), which is responsive to geosmin; the AL3 glomerulus (blue), tuned to nonanal; and the AM2 glomerulus (magenta), tuned to lilac aldehyde. (**I**) Concentration dependency of the PD3, AL3 and AM2 glomeruli to their cognate odorants (geosmin, nonanal, and lilac aldehyde, respectively). The glomeruli showed significantly different dose response curves (F_1,105_ = 21.5; p < 0.05), with the PD3 glomerulus having the lowest EC_50_ (10^−9^ concentration) compared to AL3 (10^−5^) or AM2 (10^−4^). (**J**) Tuning curve for the PD3 glomerulus to a panel of 21 odorants, each tested at 10^−2^ concentration.

In *Drosophila*, geosmin selectively activates a single class of OSNs, which in turn express a receptor exclusively tuned to this compound [Stensmyr *et al*., 2012]. Thus we wondered if *Ae. aegypti* detects geosmin with similar specificity. To address this issue, we next turned to functional imaging, using an *Ae. aegypti* knock-in strain (*PUb-GCaMP6s*) carrying pan-neural expression of the calcium sensitive reporter GCaMP6s from the *ubiquitin* locus [Bui *et al*., 2018]. *PUb-GCaMP6s* mosquitoes were glued to holders that permitted two-photon imaging of calcium responses in the antennal lobe (AL)[Vinauger *et al*., 2018](**Figure 3B, C**). Imaging across the AL revealed no significant responses to geosmin in the vast majority of glomeruli (**Figure 3D**); however, one single glomerulus, located approximately 75 μm from the ventral surface of the AL, showed strong responses to geosmin (**Figure 3C-E**). To identify and register this glomerulus and other glomerular regions-of-interest, we mapped our two-photon imaging results to an AL atlas (Ignell *et al*., 2005). Results showed that the geosmin glomerulus was the third posterodorsal glomerulus (PD3)(**Figure 3E, F**) and demonstrated strong calcium-evoked responses to this compound that were time-locked to the stimulus onset (**Figure 3G**).

To determine the sensitivity and tuning of this glomerulus, we next examined PD3 responses under a range of geosmin concentrations (10^−2^ to 10^−12^), and compared to AL3 and AM2 glomeruli, which are tuned to nonanal and lilac aldehyde, respectively (**Figure 3H**). Compared to these other glomeruli and their cognate odorants, PD3 exhibited orders of magnitude higher sensitivity to geosmin (**Figure 3I**), with an ED_50_ of 1.75×10^−9^ and strong responses at picogram levels. By contrast, the ED_50_s of AL3 and AM2 to nonanal and lilac aldehyde, respectively, were 1.02×10^−5^ and 1.92×10^−4^. When factoring in the effects of vapor pressure, these differences become even greater: geosmin has a 100- to 500-fold lower vapor pressure than nonanal and lilac aldehyde (0.001 mmHg, compared to 0.1 and 0.532 mmHg, respectively), causing the airborne concentrations of geosmin to be even lower than that of nonanal or lilac aldehyde.

Given PD3’s extreme sensitivity to geosmin, we next examined how this glomerulus responded to a panel of different odorants, including compounds important for mosquito host-detection, oviposition-site selection, and those commonly used as repellents. From this panel, geosmin elicited the greatest PD3 response, with a 2 to 20-fold higher response compared to the other odorants (**Figure 3J**). Interestingly, odorants that elicited the next greatest responses were p-cresol and hexanoic acid, odorants suggested to be involved in oviposition-site choice and blood host-selection [Knight and Corbet, 1991; Baak-Baak *et al*., 2013; Afify *et al*., 2014]. Although PD3 showed narrow tuning to geosmin, as measured by the kurtosis of the tuning curve (a measure of the peakedness of the distribution), with a value of 6.3, it lacked the tuning of the DA2 glomerulus of *Drosophila melanogaster* to geosmin, which has a kurtosis value of 16.2 [Stensmyr *et al*., 2012]. Nonetheless, PD3’s extreme sensitivity and specificity to geosmin indicates that this single olfactory circuit is biologically important for *Ae. aegypti* mosquitoes.

### Geosmin works as an oviposition attractant in the field

So far, we have demonstrated that geosmin confers oviposition site preference in the laboratory. We subsequently wondered if geosmin also works under field conditions as a potential tool to disrupt the life cycle of *Ae. aegypti*. To evaluate this approach we chose a site with high *Ae. aegypti* incidence, namely Miami (Florida, USA) where combatting mosquitoes has been a top priority since the arrival of the Zika virus in 2016 [Grubaugh *et al*., 2017]. The field study was conducted across the greater Miami area at 21 sites over the course of seven weeks (**Figure 4A**), using custom-made ovitraps (See Experimental procedures), baited with sachets containing dilutions of synthetic geosmin (20 mL of either a 10^−3^, 10^−4^ or 10^−5^ dilution)(**Figure 4B**). The geosmin-baited ovitraps with the 10^−4^ dilution held an increased number of eggs in comparison to control traps (baited with solvent only)(**Figure 4C**). Curiously, traps baited with the higher, or lower concentration did not cause an oviposition preference in comparison to water alone (**Figure 4D, E**). Geosmin accordingly works as an oviposition attractant within a narrow concentration range – a phenomenon previously observed also for other oviposition stimulants in *Aedes*, where attraction is only elicited within a narrow concentration range [Afify and Galizia, 2014]. The molecular and neuronal basis of this phenomenon remains unknown. Nevertheless, these experiments serve as proof-of-concept that geosmin can indeed function in attract-and-kill mosquito control approaches.

**Figure 4.**
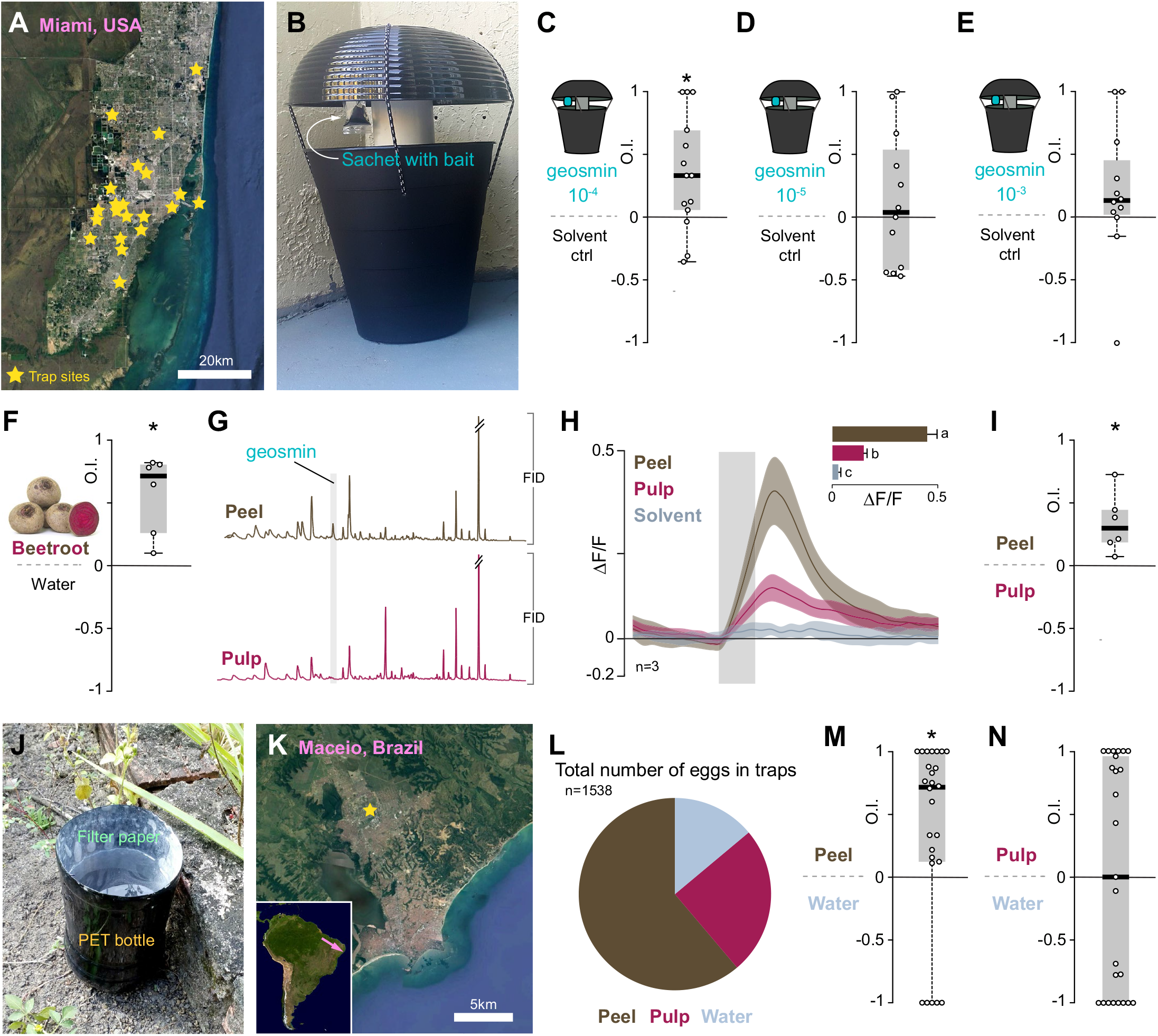
Geosmin as a mosquito control agent. (**A**) Map over greater Miami area with trap sites marked. Satellite image courtesy of Google maps. (**B**) Oviposition trap used for the field experiments. (**C-E**) Oviposition indices (OI) from Miami mosquitoes offered a choice between control traps (water only) and traps baited with geosmin. Each data point represents the average OI from a single site (n = 11-14). Box plots as per Figure 1C. Preference was tested with one-sample Wilcoxon test, theoretical mean 0. Star denotes significantly different from 0; p < 0.05. (**F**) OI of WT mosquitoes (20 mosquitoes/trial, n=6 trials) from a binary-choice test between beet extract and water. Box plots as per Figure 1C. Preference was tested with one-sample Wilcoxon test, theoretical mean 0. Star denotes significantly different from 0; p < 0.05. (**G**) Flame ionization detection (FID) traces from a gas chromatography-mass spectrometry analysis of head space comparing volatiles emitted from beetroot peel and beetroot pulp. (**H**) PD3 responses (ΔF/F_0_) to the extracts of the beet rind (brown), pulp (purple), and solvent (methanol) control (blue). Grey bar denotes the time course of odor stimulus. Traces are the mean; area is the SEM (n = 3 mosquitoes). *Inset*: Mean responses to the extracts. Letters denote significant differences between stimuli (Kruskal-Wallis test: X = 63.19, p < 0.0001; multiple comparisons: p < 0.05). (**I**) OI of WT mosquitoes (20 mosquitoes/trial, n=6 trials) from a binary-choice test between beetroot pulp extract and beetroot peel extract. Box plots as per Figure 1C. Preference was tested with one-sample Wilcoxon test, theoretical mean 0. Star denotes significantly different from 0; p < 0.05. (**J**) Simple oviposition trap constructed from painted PET bottles lined with filter paper used for the experiments in Brazil. (**K**) Brazil field site. Satellite image courtesy of Google maps (**L**) Total number of eggs in traps. (**M**) OI from wild mosquitoes offered a choice between control traps (water only) and traps baited with beetroot peel extract. Each data point represents a collection event (n=26). Box plots as per Figure 1C. Preference was tested with one-sample Wilcoxon test, theoretical mean 0. Star denotes significantly different from 0; p < 0.05. (**N**) OI from wild mosquitoes offered a choice between control traps (water only) and traps baited with beetroot pulp extract. Each data point represents a collection event (n=26). Box plots as per Figure 1C. Preference was tested with one-sample Wilcoxon test, theoretical mean 0. Star denotes significantly different from 0; p < 0.05.

### Geosmin can be substituted by beetroot juice

Unfortunately, geosmin is both expensive and difficult to obtain, particularly in the developing tropical and subtropical countries where *Ae. aegypti* is causing most harm. Thus, unless a cheap source of geosmin can be identified, our findings would be of little practical consequence. Therefore, we next set out to find a more readily available source of geosmin for use in vector control. The distinct odor of geosmin is responsible for the earthy smell of beetroots (*Beta vulgaris*)[Lu *et al*., 2003a]. Beetroots can be grown throughout much of the world and require fairly simple farming procedures. We thus wondered if beetroot juice could be used as a substitute oviposition lure. Indeed, cups spiked with extract from beetroots contained significantly more eggs than cups with water alone (**Figure 4F**). We next wondered if geosmin alone, or if also other chemicals present in beetroot, mediate the observed preference. In this context beetroots carry their own internal control; geosmin is reportedly produced and enriched in the peel, whereas the pulp only contains trace amounts of this compound [Lu *et al*., 2003a; 2003b], which we also confirmed using GC-MS (**Figure 4G**).

To examine whether beetroot evoked responses in the same olfactory channel as synthetic geosmin, we again conducted calcium imaging experiments using the *Pub-GCaMP6s* line. When stimulated with an extract of the beetroot peel, PD3 elicited strong calcium-evoked responses significantly greater than the solvent (**Figure 4H**). By contrast, an extract of the beetroot pulp elicited significantly lower responses compared to the peel, although still higher than the solvent control (**Figure 4H**). Importantly, responses to the beetroot peel were on the same order as responses to geosmin (p = 0.88)(**Figure 2G**). In line with the imaging results, gravid females also strongly preferred to lay eggs in cups with peel extracts over those containing pulp extract (**Figure 4I**). In summary, beetroot peel is a cheap and viable alternative to geosmin for use in mosquito population control.

### Beetroot juice baited traps catch mosquitoes in Brazil

Having acquired promising results with beetroots under laboratory conditions, we next conducted a small-scale field study. We performed the experiments in Northeastern Brazil (state of Alagoas), which is an impoverished region with a high incidence of mosquito transmitted infectious diseases [Heukelbach *et al*., 2016]. We first devised a simple oviposition trap, constructed from used PET bottles, painted black and lined with filter paper (**Figure 4J**), which we baited with extracts of beetroot peel or pulp, and with water only as control. We placed traps around the campus grounds of the Federal University of Alagoas in Maceió (**Figure 4K**), an urban area with a high mosquito frequency. In line with the lab results, traps baited with peel extract yielded considerably more mosquito eggs than traps with water alone (**Figure 4L, M**). Traps baited with pulp extract, however, held about as many eggs as water (**Figure 4L, N**). To identify the attracted mosquitoes, the eggs were hatched. Out of the 335 adult mosquitoes that emerged, 97.7% were *Ae. aegypti* and *Aedes albopictus*, and the remaining *Culex* spp. In short, beetroot peel works as an oviposition stimulant under field conditions in the tropics, and might accordingly be an inexpensive and environmentally friendly method for mosquito control in developing countries. The simple trap design can be improved, as can the beetroot formulation, to increase trap catches. Adapting traps to local conditions, e.g. making traps less prone to be carried away or destroyed by locals, human and non-human, as well as preventing runaway microbial activity by including anti-fungal chemicals in the formula (which can also be plant based), might be needed. Nevertheless, our findings provide an innovative method in which to use a natural lure to interrupt the life cycle of *Ae. aegypti*.

## CONCLUSION

We show here that geosmin confers preferential egg-laying in *Ae. aegypti*, which (presumably) associates this chemical with microbes, such as cyanobacteria, present in the aquatic habitats of the larvae. *Aedes* larvae likewise find geosmin attractive, as well as geosmin-producing cyanobacteria. Using *in vivo* two-photon imaging we find that adult *Ae. aegypti* detect geosmin with a high degree of sensitivity and selectivity, with geosmin solely activating a single glomerulus, innervated by sensory neurons responding to geosmin already at extremely low dilutions (10^−11^). Finally, field experiments performed in Miami and Brazil with synthetic geosmin, and geosmin derived from beetroot peel respectively, demonstrate the possibility of using geosmin as bait in trap-and-kill mosquito control approaches.

The similarity by which *D. melanogaster* and *Ae. aegypti* detects and decodes geosmin is striking. Both species are equipped with highly sensitive and selective detection machineries for this microbial volatile. The precise receptor in *Aedes* mediating the geosmin response from the PD3 glomerulus remains to be uncovered. However, since no direct ortholog of the *Drosophila* geosmin receptor *Or56a* is found in the *Ae. aegypti* genome [Matthews *et al*., 2018], it is clear that these two species have derived their superb ability to detect geosmin independently. It is intriguing that the same chemical, which appears to carry the same message, i.e. presence of microbes, induces opposing valence in these two species. How other Dipterans, or other insects for that matter, react to and decode this ubiquitous compound would certainly be interesting to determine.

Many geosmin producing microbes, including cyanobacteria, produce toxins [Carmichael, 1992]. In fact, certain strains of cyanobacteria are also acutely toxic to *Ae. aegypti* [Kivarante *et al*., 1993]. The *Kamptonema* sp. PCC 6506 strain used in this study produces the neurotoxin anatoxin-a (or Very Fast Death Factor, VFDF) as well as the cytotoxin cylindrospermopsin [Méjean *et al*., 2010; Mazmouz *et al*., 2010]. Possibly, mosquito larvae might have a certain degree of tolerance for cyanobacterial toxins, akin what is found in lake flies (Chironomidae) and shore flies (Ephydridae), which habitually feed on cyanobacterial mats [Krivosheina 2008]. Not all cyanobacteria are toxic, however, and mosquitoes might be endowed with other means, olfactory and/or gustatory, to separate harmful cyanobacteria from harmless.

Apart from offering insights into how insects and mosquitoes in particular decode odors, our findings also provide a novel and sustainable approach for mosquito control. The use of beetroot peels as bait carries the benefit that the part of the beetroot that would otherwise have gone to waste, now has its distinct use. Whereas the peel can be used to trap mosquitoes, the pulp can be used to make borscht [Blankensteen, 1974], or some other tasty and nourishing meal.

## ACKNOWLEDGEMENTS

We thank Christopher Potter for valuable comments on the manuscript, and members of our respective labs for critical feedback during the project. We thank Elisabeth Barane for expert assistance with growing cyanobacteria, and Prof Antonio Euzebio Goulart Sant’ana for support with the Brazilian fieldwork. N.M. and M.C.S was supported by the Craaford Foundation, Carl Tryggers Foundation, and the Swedish Research Council. A.L.C., R.A., and M.D. were supported by The Florida Department of Agriculture and Consumer Services (Grant name “*Highly attractive biological insecticide trap (*HABIT*) to reduce mosquito populations*”) and The Centers for Disease Control and Prevention (CDC), Southeastern Center of Excellence in Vector-borne Disease. We would like to thank the entire DeGennaro laboratory for their help in obtaining the field collections of mosquito eggs. G.H.W. and J.A.R. were supported by awards from the National Science Foundation (*IOS-1354159*), Air Force Office of Scientific Research (*FA9550-16-1-0167*), and University of Washington Innovation Award. M.G. and the Pasteur Collection of Cyanobacteria is funded by the Pasteur Institute. The content is solely the responsibility of the authors and does not necessarily represent the official views of respective employers or the CDC.

## AUTHOR CONTRIBUTIONS

N.M. and M.C.S. conceived the study. N.M. performed electrophysiology, and all lab oviposition and feeding experiments, with support from M.F.T., who also carried out the field trials in Brazil. G.H.W. and J.A.R. designed, performed, and the analyzed the two-photon imaging experiments. A.L.C., R.A., and M.D. designed and performed the field ovitrap studies in South Florida as well as provided reagents for the laboratory experiments. M.G. provided the cyanobacteria and assisted the experiments. N.M. and M.C.S wrote the manuscript, with feedback from all authors.

## EXPERIMENTAL PROCEDURES

### Mosquito rearing

*Aedes aegypti* were reared and kept in an environmental room under LD 12:12 h cycle at 26 −28 °C, 79% RH. Eggs were hatched by adding deoxygenated water with ground fish food (Tetra tabimin, 052967, Arken Zoo) inside a plastic container (L: 32 x W: 17 x H: 10 cm). Post-hatching, larvae were fed daily with ground fish food. The pupae were placed in small cups with distilled water and moved to a 30 cm^3^ mesh cage (DP100B, BugdormStore, Taiwan), and allowed to eclose. Adult mosquitoes were fed on 10% sucrose solution (weight: volume in distilled water) from a cotton wick inserted into a vial. Mosquitoes were blood-fed using an artificial blood feeder (CG-1836, Chemglass Life Sciences, USA) filled with defibrinated sheep blood (SB055, TCS Biosciences Ltd, Buckingham) (heated to 37°C), spiked with 10 mM ATP (A1852, Sigma-Aldrich) for about 2 hours per cage. Blood-fed mosquitoes were subsequently allowed to feed on 10% sucrose solution.

### Chemical reagents

Saline was made based on the Beyenbach and Masia [2002] recipe, containing 150.0 mM NaCl, 25.0 mM N-2-hydroxyethyl-piperazine-N’-2-ethanesulfonic acid (HEPES), 5.0 mM sucrose, 3.4 mM KCl, 1.8 mM NaHCO_3_ 1.7 mM CaCl_2_, and 1.0 mM MgCl_2_. The pH was adjusted to 7 with 1 M NaOH. Odorants used in calcium imaging experiments were purchased from Sigma-Aldrich or Bedoukian at the highest purity (generally >98%). Geosmin was purchased from Perfume Supply House (https://perfumersupplyhouse.com) at 1:100 concentration in dipropylene glycol (DPG) and from Pell Wall Perfumes (https://pellwall.com) at 1:10 concentration in DPG. Odorants included geosmin, terpenes: (±)linalool, lilac aldehyde (mixture of isomers), α-pinene, linalool oxide, geraniol, citronellal, geranyl acetate; aromatics: benzaldehyde, benzyl acetate, methyl benzoate, *p*-cresol, DEET; and aliphatic aldehydes, alcohols and acids: octanal, nonanal, hexenal, 1-octen-3-ol, methanol, lactic acid, and hexanoic acid. Odorants were diluted 1:100 vol vol^−1^ in mineral oil, except for geosmin and DEET, which were diluted 1:100 vol vol^−1^ in DPG and methanol, respectively.

### Oviposition assays

Oviposition assays were conducted to test four different stimuli and control (water): geosmin 10^−5^ (350 μL), beetroot peel (3 g), beetroot pulp (3 g), and cyanobacteria (250 μL). 20 blood-fed females were used per assay. 72 h post blood-feeding, two plastic containers filled with 80 mL distilled water (8 x 8 x 3 cm) were placed at opposite corners, one serving as stimulus and the other as control. Each container was lined with 5.5D Whatman filter paper (WHAT1001500, Sigma Aldrich) on the sides. Oviposition was allowed for 72 h, the number of eggs laid in each container was counted using a microscope. Number of laid egg was compared between control and treatment, and an Oviposition Index (OI) was calculated as follows: (#_treatment_ − #_control_)/(#_treatment_ + #_control_) where #_treatment_ indicates number of eggs laid in geosmin and the #_control_ indicates number of eggs laid in control.

### Capillary feeding assay

Capillary feeding assays were conducted to assess the effect of geosmin on nectar feeding behavior of female *Ae. aegypti*. This assay is adapted to mosquitoes based on similar assays for *Drosophila melanogaster* [Ja *et al*., 2007]. Starved females were transferred individually to a standard polypropylene *Drosophila* rearing vial with access to two 5 μL calibrated glass capillaries embedded in cotton plugs. One of the capillaries serves as the control, containing 10% sucrose in distilled water. The stimulus capillary contained 10% sucrose spiked with 10^−3^ geosmin (diluted from 10% Geosmin, Pell Wall Perfumes). After two hours, the remaining liquid in all capillaries was measured, by aligning a metric ruler to the tip of the capillary and measuring the height of the liquid meniscus. A vial without mosquito was included as evaporation control. A feeding index (FI) was calculated as follows: [(stimulus − evap) − (control − evap) / [(treatment − evap) + (control − evap)]. Vials were excluded if any of the mosquitoes died during the assay. No CO_2_ was added to these experiments.

### Membrane feeding assay

20 females were used per feeding assay. Two glass-jacketed membrane feeders (Chemglass Life Sciences, USA) connected through silicone tubes to a water bath (37 °C) were positioned on top of each cage. The membrane feeders were prepared by stretching a layer of laboratory parafilm (P7793, Sigma-Aldrich) over the feeders (simulating human skin), thereafter defibrinated sheep blood (SB055, TCS Biosciences Ltd, Buckingham) was transferred into each feeder. A nylon sock worn for 24 h by a human subject was placed over the parafilm in order to attract the mosquitoes. One of the membrane feeders serves as control, containing only the nylon sock. The stimulus membrane feeder contained a nylon sock with 350 μL of 10^−5^ geosmin (diluted from 10% Geosmin, Pell Wall Perfumes). A feeding index (FI) was calculated as follows: (#_stimulus_ − #_control_)/(#_stimulus_ + #_control_) where #_stimulus_. indicates the number of mosquitoes feeding on the geosmin spiked feeder and the #_control_ indicates number of mosquitoes feeding on the feeder without geosmin. Mosquitoes were counted every 5 min for 30 min. No CO_2_ was added to these experiments.

### Constrained contact assay

This assay is a modification of the arm-in-cage assay, where a human hand is exposed against the mesh on the outside of the cage. 20 non-blood fed females were allowed to probe and “try” to feed on the human hand. The stimulus was a human hand baited with 10^−3^ geosmin (diluted from 10% Geosmin, Pell Wall Perfumes) and the control was a the other hand without any added odors. Number of mosquitoes landing on the mesh touching the hand and probing were recorded continuous for 2, 4, and 6 minutes. A intended biting index (IBI) was calculated as follows: (#_stimulus_ − #_control_)/ (#_stimulus_ + #_control_) where #_stimulus_ indicates the number of mosquitoes trying to feed on the geosmin spiked hand and the #_control_. indicates number of mosquitoes trying to feed on the hand without geosmin. No CO_2_ was added to these experiments.

### Electrophysiology

Electroantennogram (EAG) recordings were performed using Ag-AgCl electrodes and glass capillaries filled with ringer solution (8,0 g L^−1^ NaCl 0,4 g L^−1^ CaCl_2_). Female *Ae. aegypti* were cold anesthetized for one minute before securing the body between sticky tape and dental wax. The glass capillary connected to the indifferent electrode was placed in the eye, whereas the glass capillary connected to the recording electrode was placed over the tip of the antennae. The signals were passed through a high impedance amplifier (IDAC-4, Syntech 2004, Hilversum, Netherlands) and analyzed using a customized software package (Syntech EAG-Pro 4.6). Ten μL aliquots of each dose of geosmin (diluted from 10% geosmin, Pell Wall Perfumes, UK) (10^−2^, 10^−3^, 10^−4^, 10^−5^, 10^−6^) was added onto a pre-cut Whatman filter paper (WHAT1001500, Sigma Aldrich) which was inserted into a sterilized Pasteur pipette. The stimuli were delivered via an air stream at a flow rate of 1 L. min^−1^ with a puff (2 s duration) at 30 s interval. Control (water) was tested at the beginning and end of each repetition. Octanoic acid (10^−3^) and 1-octen-3-ol (10^−3^), as controls, were also tested.

### Calcium imaging

Odor-evoked responses in the *Ae. aegypti* antennal lobe (AL) were imaged using the genetically-encoded *PUb-GCaMPs* mosquito line. Based on immunohostochemical studies, this mosquito line shows strong GCaMP6s expression in glia, local interneurons, and projection neurons. However, glia-like processes occurred on the exterior ‘rind’ of AL glomeruli and was restricted compared to the GFP labelling, thus enabling us to record from the central interior regions of the glomerular neuropil. A total of eighteen mosquitoes were used for all calcium experiments. Each mosquito was cooled on ice and transferred to a Peltier-cooled holder that allows the mosquito head to be fixed to a custom stage using ultraviolet glue. The stage permits the superfusion of saline to the head capsule and space for movement (Vinauger *et al*., 2018). Once the mosquito was fixed to the stage, a window in its head was cut to expose the brain, muscle and trachea were removed, and the brain was continuously superfused with physiological saline (Beyenbach and Masia, 2002). Calcium-evoked responses in the AL were imaged using the Prairie Ultima IV two-photon excitation microscope (Prairie Technologies) and Ti-Sapphire laser (Chameleon Ultra; Coherent). Experiments were performed at different depths from the ventral surface of the AL (15 to 90 μm), allowing characterization of glomerular responses to geosmin across the AL and allowing these glomeruli to be repeatedly imaged across preparations. Images were collected at 2 Hz, and for each odor stimulus images were acquired for 35 s, starting 10 s before the stimulus onset. Calcium-evoked responses are calculated as the change in fluorescence and time-stamped and synced with the stimulus pulses. After an experiment the AL was sequentially scanned at 1 μm depths from the ventral to dorsal surface to provide glomerular assignment and registration between preparations. Glomeruli were mapped and registered based on the positions and odor-evoked responses of the AL3, MD2 and AM2 glomeruli, using an AL atlas [Ignell *et al*., 2005] and the software *Reconstruct* [Fiala, 2005].

### Bacterial cultures

Two axenic strains *Kamptonema* sp. PCC 6506 and *Leptolyngbya* sp. PCC 8913 were grown in BG11 media at 22°C and 5-10 μmol photon.m^−2^.s^−1^.

### Larval assay

*Ae. aegypti* 3^rd^ and 4^th^ instar larvae were carefully removed from rearing pans, rinsed carefully with distilled water to remove any food residues, and kept in Petri dishes with distilled water for 30 min. Odorant stock was made by dissolving a specific amount of the treatment in 2% agarose. The assay was performed in a Petri dish (D: 10 x H: 1 cm) filled with distilled water. A test zone and control zone on opposite ends was determined and outlined. The odorant/control stock was placed into the dish 1 min beforehand to equilibrate, and an individual larva was gently introduced between the two zones. The water, odorant/control stock, and larvae, was changed after each repetition. Real time tracking was conducted throughout 4 min per repetition using Noldus Ethovision. Time of contact between larvae and the odorant/control zone was counted for each assay and a response index calculated as follows: (#_odorant_ − #_control_)/(#_odorant_ + #_control_) where #_odorant_ indicates time larvae spent in test zone and the #_control_ indicates time larvae spent in control zone. Respective RI values were compared with each other and analysed for statistical significance.

### Field studies

Ovitraps: The custom-made ovitrap structure (**Figure 4B**) was mounted combining 3 pieces of white polyvinyl chloride (PVC) pipes and 2 pieces made of black plastic. The body of the trap consisted of a black bucket (fniss trash can, black, item #602.954.38, IKEA, Sweden) where 4 holes were drilled at the top, closely to edge of the opened side. Two crossed rubber bungee cords were tied to the bucket by using the holes and were used to hold the rounded concave black lid (Camwear Round Ribbed Bowl, Item #:214RSB18CWBK, Cambro, CA, USA). A 40 cm cylinder-shaped 3” PVC white pipe connected at the extremities to two different PVC fittings, a bottom piece (3 in. x 3 in. x 1-1/2 in. DWV PVC Sanitary Tee Reducing, Charlotte pipe, Charlotte, NC, USA) and a top piece (3 in. white slip hub #1005, Valterra, Mexico), were used to build a central pillar which was put into the center of the bucket. The bottom PVC fitting has a lateral hole that allowed the trap to be filled up with tap water from the top of the pillar. The top piece has a squared stage for supporting the lid and also holes in each corner where the scented sachet was hanged using a metal cup hook (arrow satin nickel 7/8” cup hook, arrow™ utility hooks, Liberty Hardware Manufacturing Corporation, Winston-Salem, NC). The half-bottom of the bucket’s inside wall was coated with a round-shaped chromatography paper (Cat n# 3030-690, GE Healthcare Life Sciences, Boston, MA) as a substrate for laying. The bucket was filled up with 3 L of tap water by using the central pillar before the trap to be deployed at each site. The homemade sachets consisted of 12 cm long strips of low density polyethylene (LDPE) (2 Mil Poly Tubing Roll - 1 1/2” x 1,500’, model n. S-3521, ULINE polytubing, Pleasant Prairie, WI, USA) filled up with 20 mL of dipropylene glycol (DPG, Sigma-Aldrich^®^) for the control traps, while baited sachets were prepared with 20 mL of three different doses of Geosmin, 0.1% (10^−3^), 0.01% (10^−4^) and 0.001% (10^−5^) (diluted from 10% Geosmin, Pell Wall Perfumes, UK). The sachets were sealed by using a FS-300 hand sealer (FS-series, Sealer Sales, INC, CA, USA). and were kept individually inside of a Whirl-Pak^®^ Write-On Bags – 18 oz (product number B01065WA, Nasco, Fort Atkinson, WI) until placement in the traps.

#### Collections

The field experiments in Miami-Dade County (Miami, FL) lasted for 30 consecutive weeks over 21 sites from August 8, 2017 to March 31, 2018. For reliability purposes, we stipulated to record at least 40 positive trials (defined by the presence of at least one egg in at least one trap) for each geosmin dose tested. The 10^−4^ trials lasted for 7 weeks (from 8/11 to 10/21/2017), 10^−5^ trials for 7 weeks (from 10/28 to 12/09/2017), and 10^−3^ trials for 16 weeks (from 12/12/2017 to 03/31/2018). The custom-made ovitraps were deployed in pairs in each site. Randomly, a non-scented sachet was hung in one trap while a Geosmin-scented one was in its counterpart. The traps were setup in contact with each other (to keep similar microenvironment condition) and were exposed outside of the sites (houses, communities or apartments – no higher than the third floor) for 4 days per week. After the exposure period, the chromatography papers inside the paired traps (oviposition substrate) were collected from both control and experimental traps and were placed in respective labeled whirl pack bag until further analysis in the laboratory. The papers were qualitativly and quantitativly evaluated for mosquito egg presence under the stereo microscope (model EZ4, Leica). For species identification, the positive chromatography papers were submerged into deionized, deoxygenated water and larvae were fed with dissolved tablets of Tetramin tropical fish food (catalog #16152, Tetra, Melle, Germany). The emerged adults were identified by using morphological characters described in a morphological identification key [Consoli *et al*., 1994].

#### Maceio, Brazil

The ovitraps were installed in September 2017 at the Federal University of Alagoas, Brazil (9°33’10.6”S 35°46’30.7”W) over a period of 1 month. Modified ovitraps were made by painting PET bottles black and cutting the narrow opening, giving them the following measurements: H: 17 cm D: 9 cm. Strips of filter paper (30 cm x 5 cm) were used to line the inside of the opening of each trap. 600 mL of water was added to each trap. Two stimuli were tested: beet peel (10 g), beet pulp (10 g) against control (water). Ovitrap catches were checked and collected every 3 days and content was renewed. Collected eggs were allowed to eclose in the laboratory and adults were classified according to their morphological characteristics [Consoli *et al*., 1994].

